# A nearly 30-years living collection from the Royal Botanic Garden Edinburgh is a new species: a case study of *Betula mcallisterii* sp. nov. (sect. *Acuminatae*, Betulaceae) and its little hybridization with *Betula luminifera*

**DOI:** 10.1101/2022.10.14.512242

**Authors:** Huayu Zhang, Junyi Ding, Nian Wang

## Abstract

Species description meets challenges arising from various species concepts. Integrating different sources of information and multiple lines of evidence are important for species recognition and discovery.
Here we use morphology, restriction site-associated DNA sequencing and flow cytometry to study the species status of the unidentified *Betula* samples collected in this study and to assess the extent of hybridization between the unidentified *Betula* samples and *B. luminifera* in natural populations.
Molecular analyses show the unidentified *Betula* samples as a distinct lineage and reveal very little genetic admixture between the unidentified samples and *B. luminifera*. Unexpectedly, the unidentified samples formed a well-supported monophyletic clade with the living collection of *B. luminifera* 19933472 in the Royal Botanic Garden Edinburgh which was introduced from Southwest China. Flow cytometry shows that the unidentified samples and *B. luminifera* 19933472 are diploid.
Our data indicates that *B. luminifera* 19933472 and the unidentified *Betula* samples should be recognized as a new species, namely *B. mcallisterii*. The very little introgression between *B. mcallisterii* and *B. luminifera* indicates a strong reproductive barrier. Our research shows the importance of gathering information from wild populations and the value of an integrative approach in species discovery.

**Societal Impact Statement:** A comprehensive survey of plant species from natural populations can aid greatly in taxonomy and species delimitation. Here, we discovered a new birch species from the wild and unexpectedly found that this species has been introduced to the Royal Botanic Garden Edinburgh for nearly 30 years. We found very little introgression between the new species and its closely-related species. Our study highlighted the importance in integrating sources of information from natural populations and botanic gardens for species discovery.

## 1. INTRODUCTION

Wild populations of a species harbor valuable information for taxonomy and species delimitation, some of which are absent from herbarium specimens, such as features of bark peeling and the level of fragrance. Although living collections in botanic gardens may supplement such valuable information, their limited number may fail to capture sufficient information compared with wild population. Hence, integrating information from wild populations would greatly facilitate species identification and sometimes aid in species discovery. For example, the recently discovered *Victoria boliviana* was initially suspected to be *Victoria amazonica* and living materials from its original habitats helped to resolve its species status (Smith et al., 2022).

Most species were described based on morphological characters, which are sometimes less reliable as morphology varies considerably across the environments, resulting in a morphological continuum (Beatty et al., 2016; Wang et al., 2014a) or morphological convergence (Bickford et al., 2007; Funk et al., 2012). Further complicating species delimitation is interspecific gene flow as it may blur species boundaries and give rise to hybrids (Bardy et al., 2011). Some hybrids possess a combination of parental morphological characters and sometimes were regarded as species (Zhang et al., 2007).

To delineate species more reliably, an integrated approach has been increasingly adopted, combining morphological, molecular and (-or) cytogenetic data (Alors et al., 2016; Fujita et al., 2012; Newton et al., 2020; Wang et al., 2022). The advantage to use an integrated approach for species delimitation is to overcome the subjective species concepts (Carstens et al., 2013; Hausdorf, 2011; Hey, 2006). Morphological species concept emphasized morphological characters and biological species concept focus on reproductive isolation (Mayr, 1942). In plants, diploid and tetraploid close relatives usually have a strong reproductive isolation as triploids are mostly sterile, impeding interspecific gene flow (Fowler & Levin, 1984; Husband & Sabara, 2004; Levin, 1975; Roccaforte et al., 2015). Nowadays, an increasing number of studies use an integrated approach to delimitate taxa and describe new species (Barrett & Freudenstein, 2011; Wang et al., 2022). The rapid development of next-generation genotyping allows for characterizing new species confidently as using hundreds and thousands of single nucleotide polymorphisms (SNPs) not only allows for robustly inferring the phylogenetic position but also for confidently estimating population genetic structure and inferring ploidy level indirectly (Cariou et al., 2013; Grewe et al., 2017; Zohren et al., 2016).

*Betula* (birch) includes approximately 65 species and subspecies, which are broadly distributed across the northern hemisphere (Ashburner & McAllister, 2016). *Betula* is well known for frequent hybridization (Anamthawat-Jónsson & Tómasson, 1999; Bona et al., 2018; Ding et al., 2021; Thórsson et al., 2010; Tsuda et al., 2017; Wang et al., 2014b), due to wind-pollination, self-incompatibility and lack of complete reproductive barriers (Ashburner & McAllister, 2016). *Betula* species have substantially morphological variation. For example, leaf or bark color vary considerably among *B. pendula* populations (Tarieiev et al., 2019). In addition, *Betula* species have a series of ploidy, ranging from diploid to dodecaploid (Ashburner & McAllister, 2016; Wang et al., 2016). Consequently, *Betula* has a very tough taxonomy and species misidentification often occurs (Wang et al., 2016).

In this study, we came across some individuals, which have similar fruits and overlapped phenology with its sympatric species *Betula luminifera*. However, these individuals have peeled bark whereas *B. luminifera* has smooth bark. We termed these individuals as the “unidentified sample” hereafter. We integrated multiple lines of evidence to resolve the species status of the “unidentified sample” and then investigate if it hybridizes with *B. luminifera*. We further show that the sample labeled as *Betula luminifera* 19933472 (*B. luminifera* 19933472 hereafter) from the Royal Botanic Garden Edinburg and the “unidentified sample” refer to the same species, which is new to science.

The aims of our study are to (1) resolve the species status of the “unidentified sample”; (2) to confirm that *B. luminifera* 19933472 is the “unidentified sample”; (3) to investigate the extent of hybridization between the “unidentified sample” and *B. luminifera*. To this end, we collected 38 the “unidentified sample” and 48 *B. luminifera* samples from four and 13 natural populations, respectively. We conducted RAD sequencing on these samples and obtained key morphological characters and ploidy level information. Our study not only highlighted the application of an integrative approach for resolving species status but also highlighted the importance in combining valuable information from natural populations and botanic gardens for species recognition and discovery.

## 2. MATERIALS AND METHODS

### 2.1 Species identification and sampling

We collected *B. luminifera* and the “unidentified sample” between May and September of 2019, 2020, and 2021 from 13 and four populations, respectively (Figure 1A). Adjacent samples were separated by ~20m. *Betula luminifera* was identified based on morphological description according to Flora of China and the taxonomical monograph of *Betula* species (Ashburner & McAllister, 2016; Li & Skvortsov, 1999). Key features to recognize *B. luminifera* include a single pendulous female catkin in raceme, smooth bark, the fruiting period between April and June and a strong fragrance from fresh cambial tissues (Figure 1B). To further accurately identify *B. luminifera*, we included samples from Guizhou province where *B. luminifera* commonly occurs. We grouped individuals as the “unidentified sample”, based on the following characteristics: a single pendulous female catkin in raceme, peeled bark, the fruiting period between April and June and no obvious fragrance from fresh cambial tissues (Figure 1C). A GPS system (UniStrong) was used to record the coordinate points of each population. Detailed sampling information is provided in Table S1.

**Figure 1.**
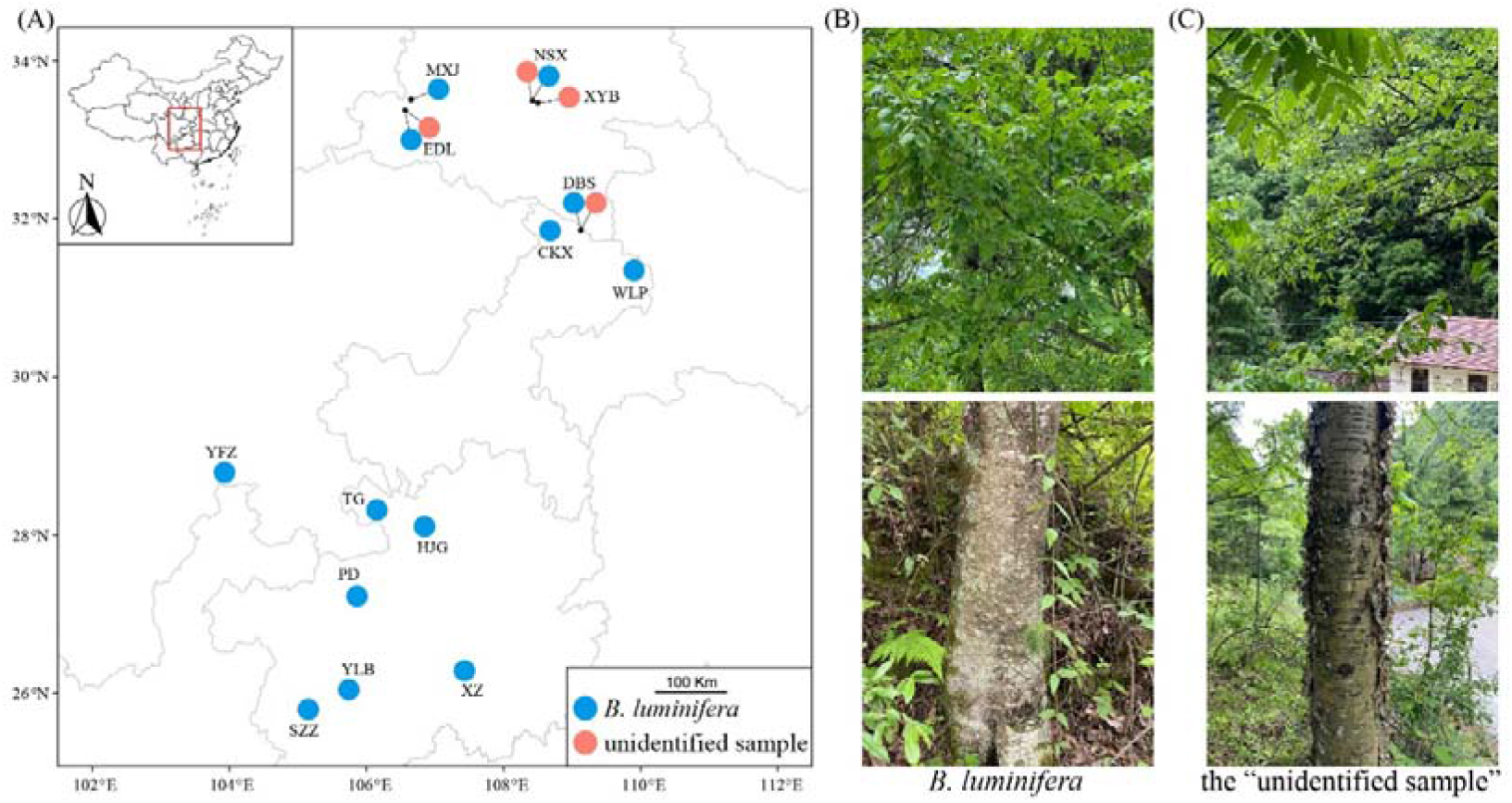
Sampling localities (A), female catkins and barks of *B. luminifera* (B) and the “unidentified sample” (C).

### 2.2 DNA extraction, sequencing and reads filtering

We extracted high quality DNA from cambial tissues following a modified 2x CTAB (cetyltrimethylammonium bromide) protocol (Wang et al., 2013). Extracted DNA was assessed with 1.0% agarose gels.

### 2.3 ITS sequencing

We amplified nuclear ribosomal (nr)ITS for 25 individuals of the “unidentified sample” using primers ITS4 (White et al., 1990) and ITSLeu (Baum et al., 1998). Reactions were performed following Hu et al. (2019). PCR products were sequenced at Tsingke Company (Qingdao, China). ITS sequences were deposited at NCBI with GenBank accession numbers OP263695-OP263719.

### 2.4 RADseq and reads filtering

A total of 86 DNA samples were selected for RADseq using an Illumina HiSeq 2500 and 150-bp pair-end sequencing with the restriction enzyme PstI (Personalbio company, Shanghai, China). RADseq data of four samples of section *Acuminatae* in Wang et al. (2021) were included for population genomic analyses, representing one each of *B. alnoides, B. cylindrostachya, B. hainanensis* and *B. luminifera* 19933472. Raw data were trimmed using Trimmomatic (Bolger et al., 2014) in paired-end mode. Reads with a quality of below 20 within the sliding-window of 5 bp and unpaired reads were discarded. Then a SLIDINGWINDOW step was performed to discard reads shorter than 40 bp.

### 2.5 Reads mapping and SNP calling

Filtered reads of each sample were mapped to the whole genome sequence of *B. pendula* (Salojärvi et al., 2017) using BWA-MEM v.0.7.17-r1188 algorithm in BWA with default parameters (Li & Durbin, 2009). Alignments were converted to sorted and indexed bam files using SAMtools v1.8 (Li et al., 2009). The MarkDuplicates tool and HaplotypeCaller from GATK v 4.1.4 were used to mark duplicates and call genotypes for each sample, respectively (DePristo et al., 2011; McKenna et al., 2010). The GenomicsDBImport was used to merge gVCF files into a combined VCF file, which used for joint genotyping using the GenotypeGVCFs tool. SNPs were filtered using a mapping quality (MQ) threshold of 40, a variant confidence (QUAL) of 30, a normalized QUAL score (QD) of 2.0, a maximum symmetric odds ratio (SOR) of 3.0, a minimum depth (DP) of 5 and a maximum depth of 200, a maximum probability of strand bias (FS) of 60.0, an excess heterozygosity (ExcessHet) of 54.69, a minimum Z-score of read mapping qualities (MQRankSum) of -12.5 and position bias (ReadPosRankSum) of −8.0, respectively. The SelectVariants filtering tool was applied to select SNPs present in at least xx of the samples. BCFtools v1.8 was used to remove SNPs within a 50 kb window with r^2^ > 0.5 to reduce linkage disequilibrium and then was used to remove SNPs with a minimum allele frequency (MAF) <=0.02 (Li, 2011). All sequences were deposited in the NCBI-Sequence Read Archive (SRA) repository under the BioProjectID PRJNA871086.

### 2.6 Population genomic analyses

A principal component analysis (PCA) was performed on SNPs of *B. luminifera*, the “unidentified sample” and the four samples of section *Acuminatae* using the ‘adegenet’ R package 2.1.1 (Jombart, 2008). The SNPs were also analyzed in ADMIXTURE v1.3.0 (Alexander & Lange, 2011), setting K from one to ten with 20 replicates for each K value. Cross-validation error estimation was performed in order to assess the most suitable value of K (Alexander & Lange, 2011). Replicate runs were aligned and visualized in pong v1.4.9 with the greedy algorithm (Behr et al., 2016).

### 2.7 Inference of ploidy level based on flow cytometry and SNPs

We performed flow cytometry following Wang et al. (2022) for three accessions of the “unidentified sample” collected from populations NSX and XYB. We also inferred the ploidy of the 86 samples sequenced in this study using the method described in Zohren et al. (2016). Briefly, we plotted the distribution of allele ratios from read counts at heterozygous sites and we would expect a peak around 0.50 for a diploid, peaks around 0.33 and 0.67 for a triploid and peaks close to 0.25, 0.50 and 0.75 for a tetraploid.

### 2.8 Phylogenetic analyses based on ITS and SNPs

Seventy additional ITS sequences from Betulaceae (Wang et al., 2022; Wang et al., 2016) were included to infer the phylogenetic position of the “unidentified sample”. A total of 95 sequences were aligned using BioEdit v.7.0.9.0 (Hall, 1999) with default parameters.

We included RADseq data of 23 *Betula* taxa from a previous study (Wang et al., 2021) and of 43 samples generated in the present study for phylogenomic analysis. *Alnus inokumae* was selected as the outgroup. SNPs were concatenated into a supermatrix with missing data below 50%. This resulted in 2,291,159 SNPs.

We analysed the ITS alignment and the supermatrix of SNPs separately, with a rapid bootstrap analysis under a GTR + GAMMA nucleotide substitution model in RAxML v.8.1.16 (Stamatakis, 2006). A total of 100 bootstraps and ten searches using the ML were performed.

## 3. RESULTS

### 3.1 Read mapping and variant calling

The individual read mappings of *B. luminifera* and the “unidentified sample” from the present study resulted in 8.8% to 95.1% of mapped reads per individual. After filtering, 340,979 variants were present in at least one individual (Table S1)

### 3.2 Population structure

PCA results based on the 40,209 biallelic SNPs revealed a clear separation between *B. luminifera* and the “unidentified sample” on PC1. *Betula* alnoides and *B. hainanensis* were separated from *B. luminifera* on PC2. *Betula cylindrostachya* formed a cluster with *B. luminifera*. The putative hybrid between *B. luminifera* and the “unidentified sample” was positioned between *B. luminifera* and the “unidentified sample”. The included *B. luminifera* 19933472 formed a cluster with the “unidentified sample”. PC1 and PC2 explained 30.7% and 3.2% of the total variation, respectively (Figure 2A).

**Figure 2.**
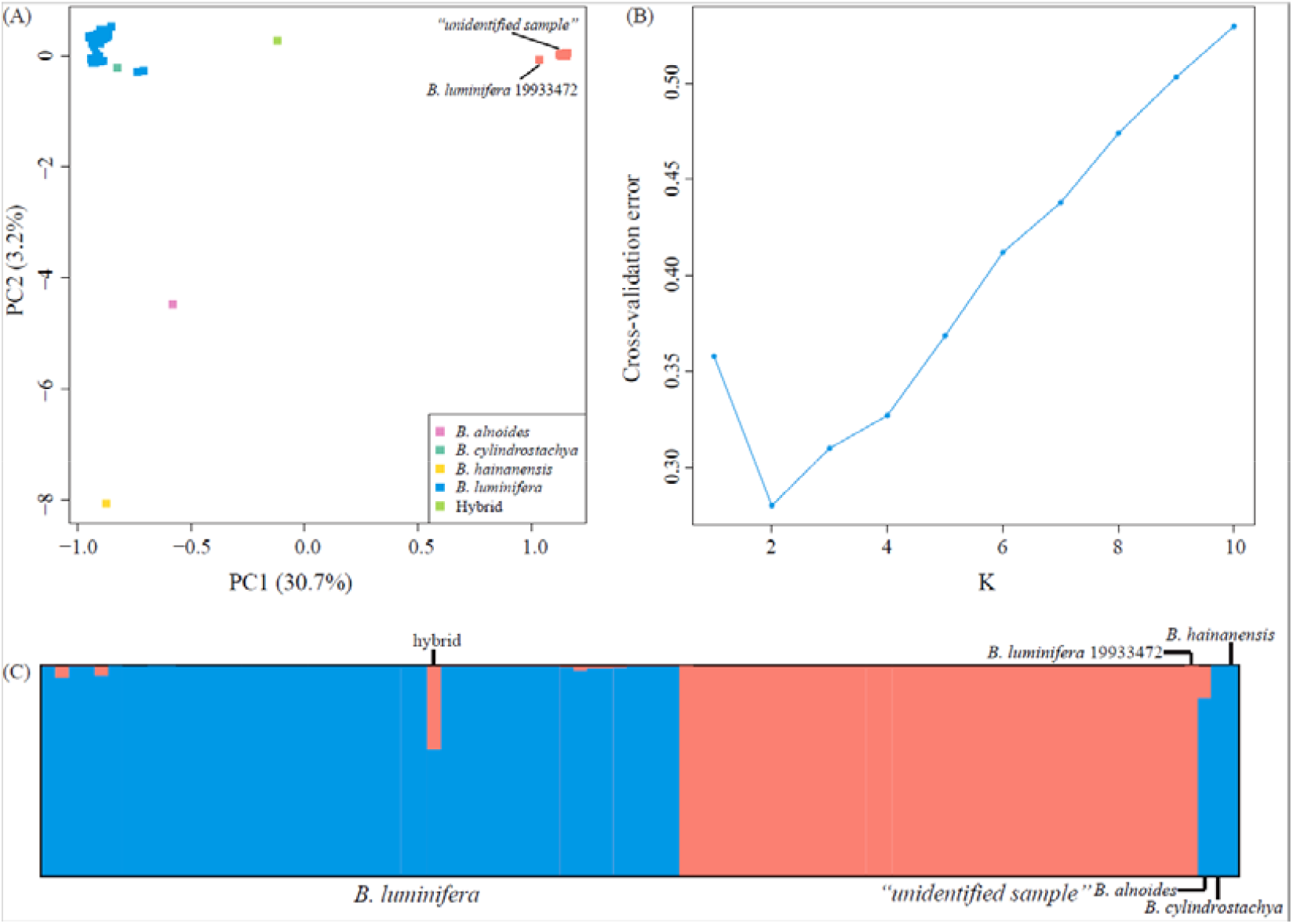
Population structure analyses based on RADseq data. (A) Principal component analysis (PCA) of *Betula* species of section Acuminate at 40,209 SNPs. (B) Cross-validation error calculated for Ks from 1 to 10. (C) Estimated genetic admixture of 90 *Betula* samples at 40,209 SNPs. Each individual is represented by a vertical line and populations are separated by a black vertical line. Blue and orange represent *B. luminifera* and *B. mcallisterii*, respectively. Previously sequenced samples (*B. luminifera* 19933472, *B. alnoides, B. cylindrostachya and B. hainanensis*) were placed on the right side.

Consistent with PCA results, admixture analyses showed the optimal K value of two (Figure 2B), corresponding to *B. luminifera* and the “unidentified sample”/*B. luminifera* 19933472. The “unidentified sample”/*B. luminifera* 19933472 formed a distinct lineage even when increased K value to six (Figure S1). Very little introgression was detected the “unidentified sample” into *B. luminifera*, with the highest admixture value of 5.31% (excluding the putative hybrid). The putative hybrid showed 39.76% and 60.24% of genetic admixture from the “unidentified sample”, and *B. luminifera*, respectively (Figure 2C).

A high level of genetic differentiation was observed between *B. luminifera* and the “unidentified sample” with F_ST_ values among populations with at least four individuals ranging from 0.30 to 0.43. Within species, F_ST_ values range from 0.01 to 0.04 for *B. luminifera*. F_ST_ value between the two populations of the “unidentified sample” is 0.02.

### 3.3 Ploidy level analyses

Flow cytometry analyses showed that the genome size of the three “unidentified sample” ranged from 388 to 445M, indicating that it is diploid (Figure S2). Plot of allele ratios at heterozygous sites showed that the “unidentified sample” has a peak around 0.50, *B. luminifera* has peaks near 0.25, 0.50 and 0.75 and the putative hybrid between the “unidentified sample” and *B. luminifera* has peaks close to 0.33 and 0.67 (Figure 3, Figures S3-4). One *B. luminifera* individual (YLB001) has ploidy level remain unclear due to the low number of reads (Figures S3).

**Figure 3.**
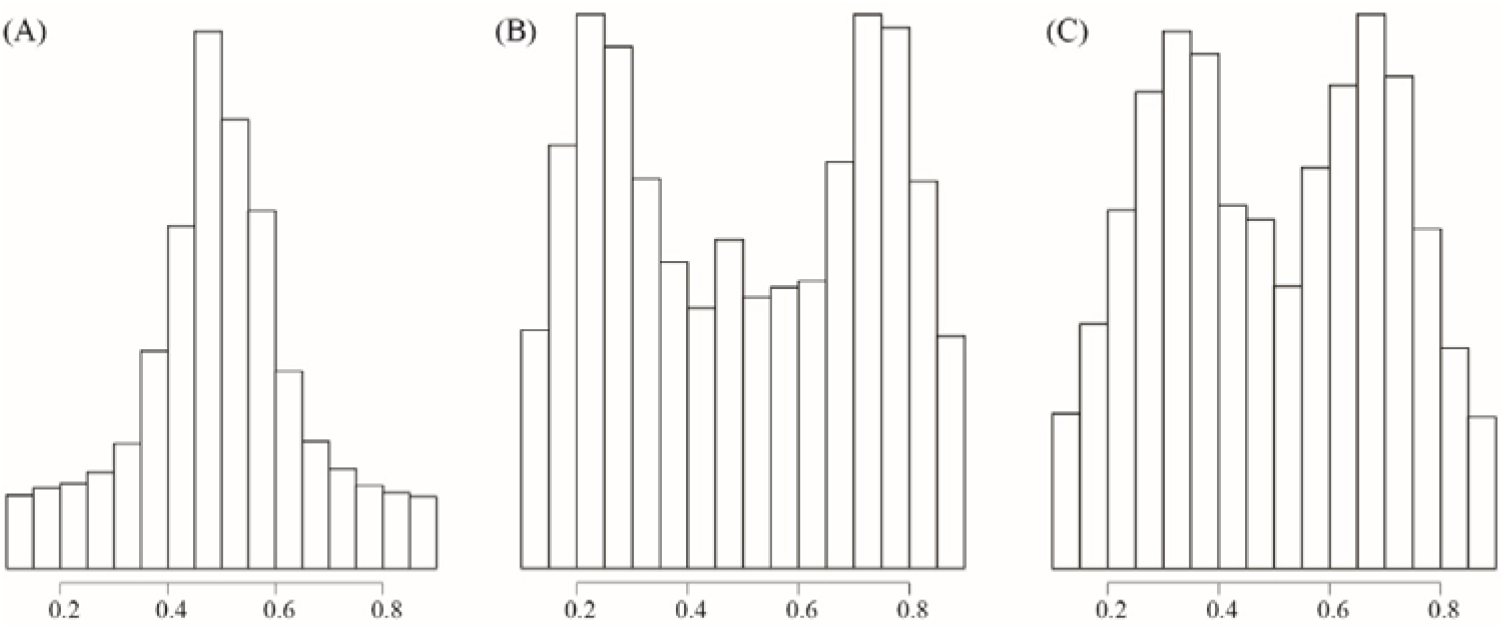
Distribution of read count ratios for heterozygous sites covered by at least 30 reads. (A) All the “unidentified sample” individuals. (B) All *B. luminifera* individuals. (C) Sample number EDL20-022, which is a hybrid between *B. luminifera* and the “unidentified sample”.

### 3.4 Identification of *Betula mcallisterii* sp. nov

We confidently establish the “unidentified sample” as a new species, namely *Betula mcallisterii*, based on morphological characters, molecular data and ploidy. The phylogenetic tree based on ITS showed that the *B. mcallisterii* tended to form a cluster except for sample EDL20-022, which was inferred as an F_1_ hybrid between *B. mcallisterii* and *B. luminifera* (Figure S5). The phylogenetic tree based on a supermatrix of 2,291,159 SNPs revealed that the *B. luminifera* 19933472 and *B. mcallisterii* formed a well-supported monophyletic clade, which is a sister to a clade of *B. luminifera, B. cylindrostachya, B. alonides* and *B. hainanensis* (Figure 4). *Betula cylindrostachya* was nested into a well-supported clade of *B. luminifera* samples (Figure 4).

**Figure 4.**
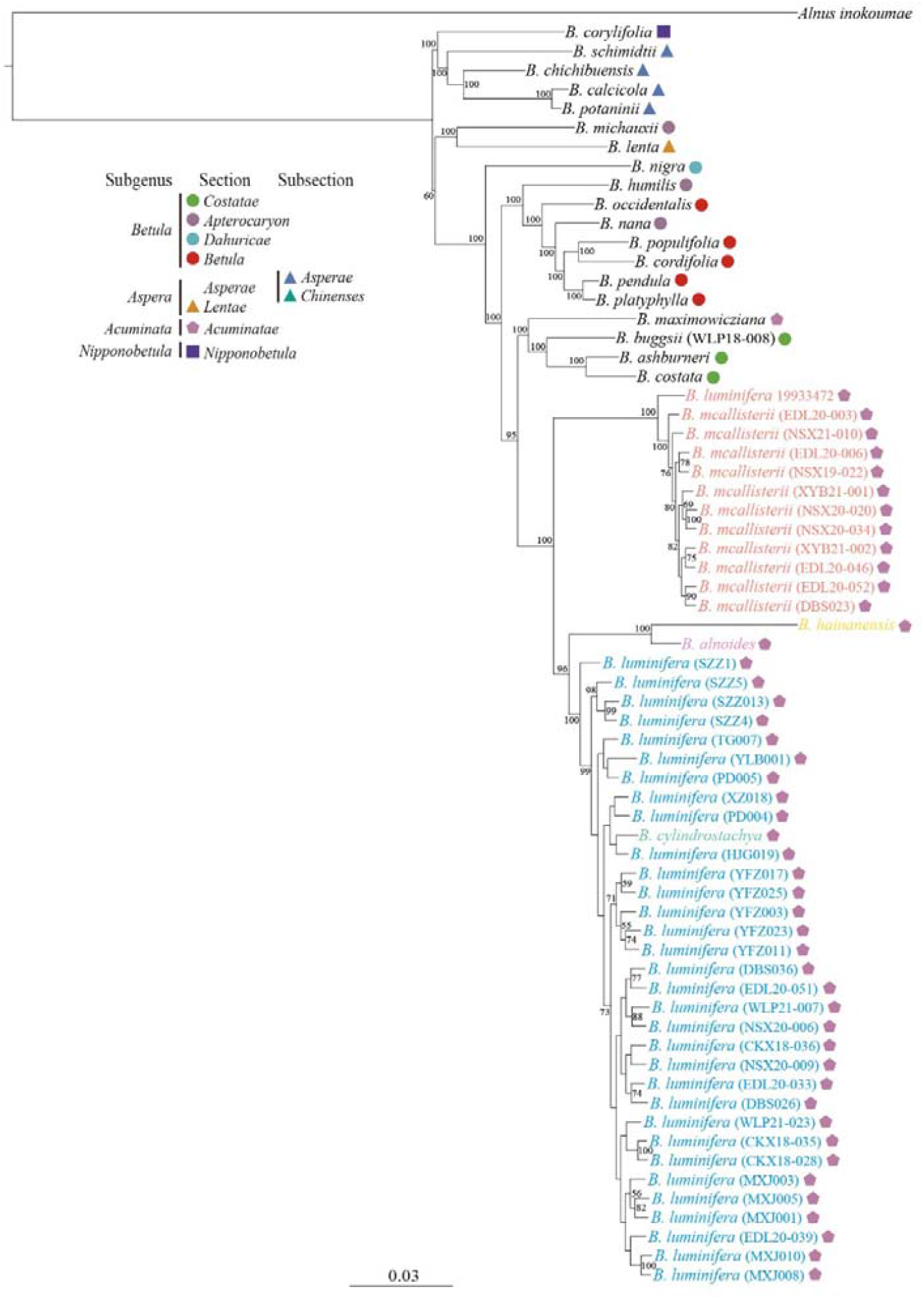
Phylogenetic tree from the maximum likelihood analysis of *B. mcallisterii* based on a supermatrix of 2,291,159 SNPs. The scale bars below indicate the mean number of nucleotide substitutions per site. Species were classified according to Ashburner, McAllister (2016). Values close to branches are bootstrap percentages of >50 %.

### 3.5 Confirmation of the *B. luminifera* 19933472 as *B. mcallisterii*

The genome size of *B. luminifera* 19933472 was 1.0 pg (Wang et al., 2016). Morphological features of *B. luminifera* 19933472 include a single pendulous female catkin in raceme and peeling bark (Figure S6). Furthermore, there is no fragrance from fresh cambial tissues (judged by five people from the RBGE).

### 3.6 Taxonomic treatment of *B. mcallisterii*

*Betula mcallisterii* Nian, sp. nov (Figure 5).

**Figure 5.**
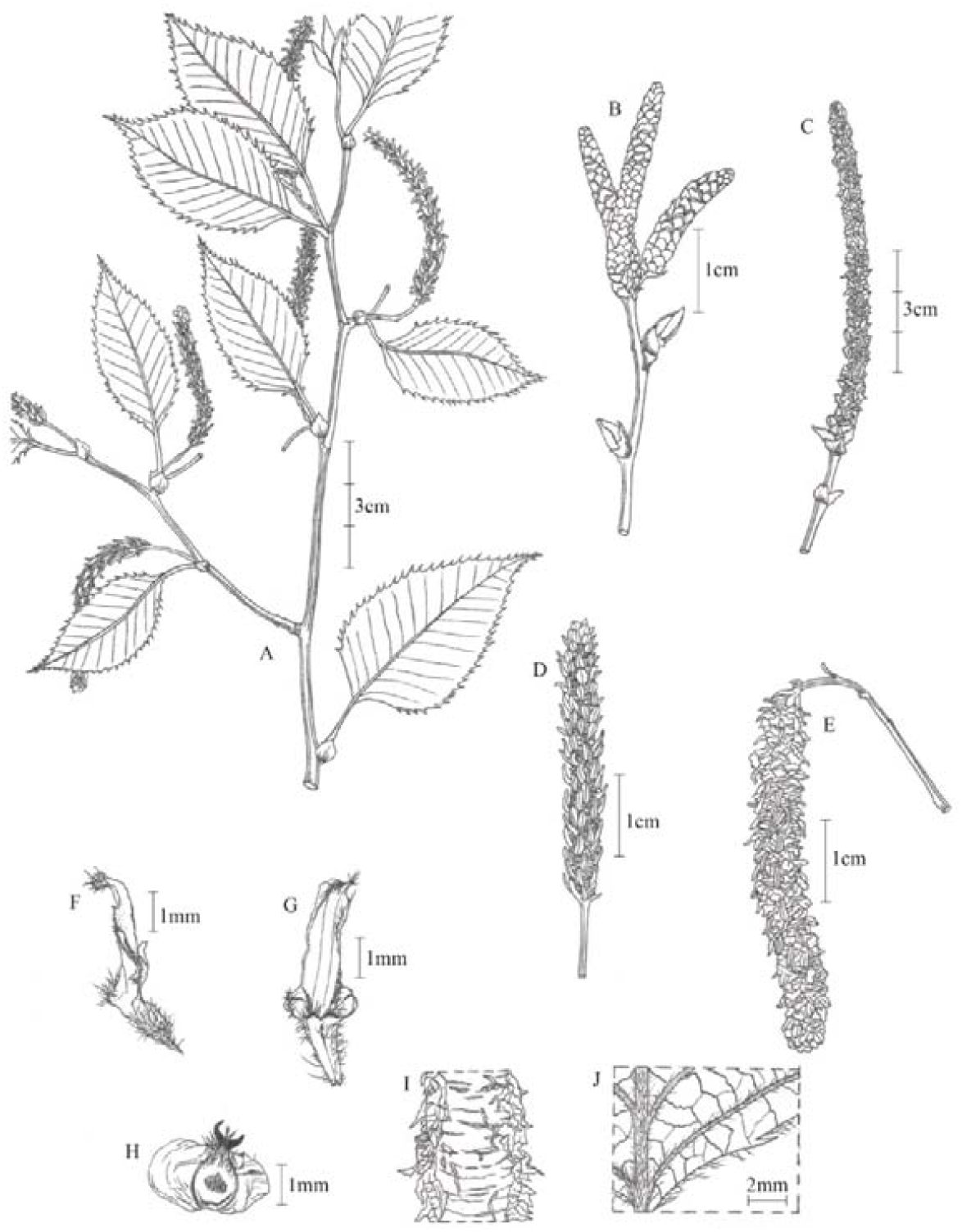
*Betula mcallisterii*. A, specimen with female catkins; B, male catkins, immature; C, male catkins, mature; D, female catkin, imature; E, female catkin, mature; F, side view of fruiting catkin scale; G, fruiting catkin scale, outer face; H, mature seed; I, bark; J, adaxial surface of leaf blade.

#### Type

CHINA. Shaanxi: Mian County, elev. ca. 1600-2000 m, 106.6 E, 33.4 N, 6 May 2021 (holotype SDAU180901; isotypes SDAU180912, SDAU180916).

#### Description

Single trunked tree to 30 m high. Trunk up to 60 cm in diameter; bark red-brown, peeling along transverse lenticels into persistent and tattered sheets. Male catkins in clusters of 3–4, pendent when matured, ~10 × 0.3 cm, length/breadth ratio ~30 : 1. Inmatured female catkins green, born singly, pendent, up to 5 cm long and 6.5 mm broad including the scales. Fruiting catkins, borne singly, long cylindrical, up to ~100 × 8 mm at maturity; Scales ~4 × 1.5 mm, margin ciliate, surfaces more or less glabrous; central lobe narrowly oblong, ~2.8 × 0.8 mm; lateral lobes ~0.5 mm wide. Seed with nutlet 2.5 × 1 mm, wing c. 1.5 mm broad each side, translucent, styles ~0.7 mm; seed surface hairy from middle toward the style.

#### Distribution and habitat

*Betula mcallisterii* occurs in the Qinling-Daba Mountains in central China from our field survey and in Yunnan province, SW China based on the collection information from *B. luminifera* 19933472. *Betula* mcallisterii grows sympatrically with *B. luminifera* at an altitude of between 1400 and 2100 meters. *Betula mcallisterii* and *B. luminifera* have no altitudinal separation. We found abundant *B. mcallisterii* individuals in EDL and NSX populations, with 16 and 30 samples confirmed as *B. mcallisterii*, respectively.

#### Etymology

*Betula mcallisterii* is named after Hugh A. McAllister, a taxonomist from Institute of Integrative Biology, University of Liverpool, for his devotion to research on taxonomy of the genus *Betula*. The Chinese name of *B. mcallisterii* is “陕南桦” (sh⍰n nán huà).

## 4. DISCUSSION

In this study, we describe *B. mcallisterii* as a new *Betula* species of section *Acuminatae* based on multiple lines of evidence. We show that *B. luminifera* 19933472 is actually *B. mcallisterii*. Furthermore, we detected very little introgression between *B. mcallisterii* and *B. luminifera* in natural populations, likely due to a difference in ploidy level. Here we discuss this new species based on an integrative approach and its natural hybridization with *B. luminifera*. We finish with the importance in integrating information from wild populations and living collections from botanic gardens in species discovery.

### 4.1 *Betula mcallisterii*, a new species of section *Acuminatae*

*Betula* mcallisterii is new species of section *Acuminatae* integrating morphological characters, molecular evidence and ploidy level information. Morphologically, *B. mcallisterii* has a single pendulous female catkin and this can be distinguished from several other species of section *Acuminatae* which have three to five pendulous female catkins in a raceme, such as *B. alnoides, B. fujianensis* and *B. hainanensis* (Zeng et al., 2008; Zeng et al., 2014). In addition, *B. mcallisterii* is different from *B. luminifera* in having obviously peeled bark (Figure 1BC). Molecularly, *B. mcallisterii* forms a distinct genetic cluster based on admixture results at the value of K=2 and onwards (Figure S1). Phylogenomic analyses strongly support *B. mcallisterii* as a monophyletic clade, separating it from other species of section Acuminate (Figure 4). Furthermore, *B. mcallisterii* fits the biological species concept which focuses on reproductive isolation (Mayr, 1942). Although we did not conduct pollination experiments, a difference in ploidy indirectly indicates reproductive barriers, especially between diploid and tetraploid (Borges et al., 2012). *Betula mcallisterii* is diploid and several other species, such as *B. alnoides, B. cylindrostachya* and *B. luminifera* are tetraploid based on genome size estimation or inference from reads covering heterozygous SNPs (Figures S2-4). A strong reproductive isolation between *B. mcallisterii* and other tetraploid species of section *Acuminatae* would be expected.

We showed that *B. luminifera* 19933472 is *B. mcallisterii. Betula luminifera* 19933472 is a diploid species (Wang et al., 2016), consistent with the ploidy level of *B. mcallisterii*. Morphologically, *B. luminifera* 19933472 has peeled bark and a single pendulous female catkin, similar with characters of *B. mcallisterii* from natural populations (Figure S6). Furthermore, phylogenomic analysis showed that *B. luminifera* 19933472 formed a monophyletic clade with *B. mcallisterii* individuals included in our present study. *Betula luminifera* 19933472 has no obvious fragrance from cambial tissues, consistent with *B. mcallisterii*. Interestingly, *B. luminifera* 19933472 was collected from Yunnan province in southwest China and the four wild populations of *B. mcallisterii* were from the Qinling-Daba Mountains. This indicates that *B. mcallisterii* has a wider distribution.

### 4.2 Hybridization between *B. mcallisterii* and *B. luminifera*

Allele sharing among closely related species is usually ascribed to incomplete lineage sorting and/-or introgressive hybridization (Twyford & Ennos, 2012). Incomplete lineage sorting is an unlikely explanation for the pattern of allele sharing observed between *B. mcallisterii* and *B. luminifera*. In most allopatric populations of *B. luminifera*, no allele transfer from *B. mcallisterii* to *B. luminifera* was detected whereas in sympatric populations either hybrid or a low level of genetic admixture was detected (Figure 2C). Such a geographic signal would not be expected from incomplete lineage sorting (Barton, 2001).

The presence of F_1_ hybrid (labeled as EDL20-022) in sympatric population EDL indicates that *B. mcallisterii* hybridized with *B. luminifera* fitting with our expectation that interploidy hybridization could occur among *Betula* species (Hu et al., 2019; Tsuda et al., 2017; Zohren et al., 2016). However, introgression between *B. mcallisterii* and *B. luminifera* was not detected in this population. In contrast, in sympatric population NSX, hybrids between *B. mcallisterii* and *B. luminifera* were not detected but trace of introgression (between 0.3% and 2%) from *B. mcallisterii* to *B. luminifera* was detected (Figure 3C). Such a difference in patterns of introgression between population EDL and NSX may reflect different histories of habitat disturbance. The area where population NSX is collected once undergone serve tree logging before China introduced Natural Forest Protection Programme and the Slope Land Conversion Programme in 1998 (Delang & Wang, 2013). Since then, the area was under strict protection and logging was totally prohibited. Different from NSX population, the area where EDL population was located is near a village and was cut through by a road allowing for car travelling. The F_1_ hybrid was collected just from roadside. The presence of F_1_ hybrid in EDL and potential ongoing habitat disturbance may have an impact on the level of genetic admixture in the future.

Similar with other studies showing asymmetric introgression from diploid to tetraploid (Clark et al., 2015; Eidesen et al., 2015; Tsuda et al., 2017; Wang et al., 2014b; Zohren et al., 2016), we observed unidirectional introgression from *B. mcallisterii* to *B. luminifera*. However, introgression from *B. mcallisterii* to *B. luminifera* is very little with the highest admixture value of 5.3%, much lower than the level of introgression from *B. nana* to *B. pubescens*, which shows the highest admixture value of 16.9% (Zohren et al., 2016). One possible explanation is that triploids between *B. mcallisterii* and *B. luminifera* may produce less viable gametes than triploids between *B. nana* and *B. pubescens*, which were reported to produce a small proportion of viable gametes (Anamthawat-Jónsson et al., 2021). Alternatively, we may have failed to sample the introgressed individuals. The fact that we did not detect hybrid swarms indicates a strong reproductive barrier between *B. mcallsterii* and *B. luminifera*, likely due to a difference in ploidy.

### 4.3 Taxonomic issues in section *Acuminatae*

Species of *Acuminatae* are characterized by the cylindrical and clustered fruiting catkins and the nutlet wings much wider than the nutlet (Ashburner & McAllister, 2016; Skvortsov, 1997). Despite the distinct morphological features, taxonomic issues exist. For example, *B. maximowiziana* (section *Acuminatae*) formed a clade with species of section *Costatae* (Wang et al., 2021). *Betula alnoides*, which was described as diploid, are found to be tetraploid based on chromosome counting of samples from several populations (pers. comm. with Hugh McAllister). Similarly, *B. luminifera*, which was described as diploid, is actually tetraploid based on our analyses. Type specimens of *Betula luminifera* were initially collected from Chenkou County with no coordinates. Though we have no idea of the exact locality of the type specimen of *B. luminifera*, samples from Chenkou County are tetraploid. Hence, the description of *B. luminifera* as diploid is likely incorrect or *B. luminifera* may have different cytotypes, like other *Betula* species, such as *B. chinensis* (Ashburner & McAllister, 2016). Further research is needed to test these hypotheses.

We found that the number of female catkins being one or two is not a reliable character for species delimitation in section *Acuminatae*. For example, *B. cylindrostachya*, a species with two female catkins in a raceme, was nested into a clade of *B. luminifera*. This is consistent with Skovotsov’s view that *B. cylindrostachya* is most closely-related to *B. luminifera* (see Ashburner, McAllister (2016)). Indeed, in natural populations, *B. luminifera* sporadically has two female catkins in a raceme (pers. comm. with Jie Zeng) and *B. cylindrostachy* occasionally has a single female catkin (Ashburner & McAllister, 2016). Further research including more *B. cylindrostachya* samples can be used to test our hypothesis that *B. cylindrostachya* and *B. luminifera* refer to the same species.

### 4.4 The role of botanic garden in species discovery and its limitation

Our journey in discovering *B. mcallisterii* shows a nice case that botanic gardens may grow undiscovered species. Interestingly, the *B. mcallisterii* has grown in RBGE for nearly 30 years before its discovery. Such cases may exist for other plant species with poor taxonomy. Our study also suggests that a clear knowledge of sufficient morphological variation from wild populations is important in species delimitation. In this study, the pattern of bark peeling, ploidy and the level of fragrance can be used to distinguish *B. mcallisterii* from *B. luminifera*. However, such information is usually unavailable or difficult to obtain from herbarium specimens. In contrast, important characters from herbarium specimens for taxonomy, such as fruits, cannot distinguish *B. mcallisterii* from *B. luminifera*. Consequently, a subset of specimens labeled as *B. luminifera* is likely misidentified.

## ACKNOWLEDGEMENTS

We thank Peter Brownless from the Royal Botanic Garden Edinburgh for sharing pictures of *B. luminifera* 19933472 and Professor Richard Buggs from the Royal Botanic Gardens Kew for valuable comments on the manuscript. The authors declare no conflict of interest related to this work. This work was funded by the National Natural Science Foundation of China (31770230 and 31600295), the Program for Introduction and Cultivation of Young Scholars in Universities in Shandong Province and the Young Scholars selected by the State Forestry and Grassland Administration, China.

## AUTHOR CONTRIBUTION

NW conceived the project. NW and JD collected samples. HZ carried out lab work. HZ and JD analyzed the data. NW and HZ wrote the initial draft.

## REFERENCES

Alexander, D. H., & Lange, K. (2011). Enhancements to the ADMIXTURE algorithm for individual ancestry estimation. BMC Bioinformatics, 12, 1–6. https://doi.org/10.1186/1471-2105-12-246

Alors, D., Lumbsch, H. T., Divakar, P. K., Leavitt, S. D., & Crespo, A. (2016). An integrative approach for understanding diversity in the Punctelia rudecta species complex (Parmeliaceae, Ascomycota). PLoS One, 11, e0146537. http://doi.org/10.1371/journal.pone.0146537

Anamthawat-Jónsson, K., Karlsdóttir, L., & Jóhannsson, M. (2021). Naturally occurring triploid birch hybrids from woodlands in Iceland are partially fertile. New Forests, 52, 659–678. https://doi.org/10.1007/s11056-020-09816-z

Anamthawat-Jónsson, K., & Tómasson, T. (1999). High frequency of triploid birch hybrid by Betula nana seed parent. Hereditas, 130, 191–193. https://doi.org/10.1111/j.1601-5223.1999.00191.x

Ashburner, K., & McAllister, H. A. (2016). The genus Betula: a taxonomic revision of birches. London, UK: Kew publishing.

Bardy, K. E., Schönswetter, P., Schneeweiss, G. M., Fischer, M. A., & Albach, D. C. (2011). Extensive gene flow blurs species boundaries among Veronica barrelieri, V. orchidea and V. spicata (Plantaginaceae) in southeastern Europe. Taxon, 60, 108–121. https://doi.org/10.1002/tax.601010

Barrett, C. F., & Freudenstein, J. V. (2011). An integrative approach to delimiting species in a rare but widespread mycoheterotrophic orchid. Molecular Ecology, 20, 2771–2786. https://doi.org/10.1111/j.1365-294X.2011.05124.x

Barton, N. H. (2001). The role of hybridization in evolution. Molecular Ecology, 10, 551–568. https://doi.org/10.1046/j.1365-294x.2001.01216.x

Baum, D. A., Small, R. L., & Wendel, J. F. (1998). Biogeography and floral evolution of Baobabs (Adansonia, Bombacaceae) as inferred from multiple data sets. Systematic Biology, 47, 181–207. https://doi.org/10.1080/106351598260879

Beatty, G. E., Montgomery, W. I., Spaans, F., Tosh, D. G., & Provan, J. (2016). Pure species in a continuum of genetic and morphological variation: sympatric oaks at the edge of their range. Annals of Botany, 117, 541–549. https://doi.org/10.1093/aob/mcw002

Behr, A. A., Liu, K. Z., Liu-Fang, G., Nakka, P., & Ramachandran, S. (2016). pong: fast analysis and visualization of latent clusters in population genetic data. Bioinformatics, 32, 2817–2823. https://doi.org/10.1093/bioinformatics/btw327

Bickford, D., Lohman, D. J., Sodhi, N. S., Ng, P. K., Meier, R., Winker, K., Ingram, K. K., & Das, I. (2007). Cryptic species as a window on diversity and conservation. Trends in Ecology & Evolution, 22, 148–155. https://doi.org/10.1016/j.tree.2006.11.004

Bolger, A. M., Lohse, M., & Usadel, B. (2014). Trimmomatic: a flexible trimmer for Illumina sequence data. Bioinformatics, 30, 2114–2120. https://doi.org/10.1093/bioinformatics/btu170

Bona, A., Petrova, G., & Jadwiszczak, K. A. (2018). Unfavourable habitat conditions can facilitate hybridisation between the endangered Betula humilis and its widespread relatives B. pendula and B. pubescens. Plant Ecology & Diversity, 11, 295–306. https://doi.org/10.1080/17550874.2018.1518497

Borges, L. A., Souza, L. G. R., Guerra, M., Machado, I. C., Lewis, G. P., & Lopes, A. V. (2012). Reproductive isolation between diploid and tetraploid cytotypes of Libidibia ferrea (= Caesalpinia ferrea)(Leguminosae): ecological and taxonomic implications. Plant Systematics and Evolution, 298, 1371–1381. https://doi.org/10.1007/s00606-012-0643-3

Cariou, M., Duret, L., & Charlat, S. (2013). Is RAD-seq suitable for phylogenetic inference? An in silico assessment and optimization. Ecology and Evolution, 3, 846–852. https://doi.org/10.1002/ece3.512

Carstens, B. C., Pelletier, T. A., Reid, N. M., & Satler, J. D. (2013). How to fail at species delimitation. Molecular Ecology, 22, 4369–4383. https://doi.org/10.1111/mec.12413

Clark, L. V., Stewart, J. R., Nishiwaki, A., Toma, Y., Kjeldsen, J. B., Jørgensen, U., et al. (2015). Genetic structure of Miscanthus sinensis and Miscanthus sacchariflorus in Japan indicates a gradient of bidirectional but asymmetric introgression. Journal of Experimental Botany, 66, 4213–4225. https://doi.org/10.1093/jxb/eru511

Delang, C., & Wang, W. (2013). Chinese forest policy reforms after 1998: the case of the natural forest protection program and the slope land conversion program. International Forestry Review, 15, 290–304. https://doi.org/10.1505/146554813807700128

DePristo, M. A., Banks, E., Poplin, R., Garimella, K. V., Maguire, J. R., Hartl, C., et al. (2011). A framework for variation discovery and genotyping using next-generation DNA sequencing data. Nature Genetics, 43, 491–498. https://doi.org/10.1038/ng.806

Ding, J., Hua, D., Borrell, J. S., Buggs, R. J., Wang, L., Wang, F., Li, Z., & Wang, N. (2021). Introgression between Betula tianshanica and Betula microphylla and its implications for conservation. Plants, People, Planet, 3, 363–374. https://doi.org/10.1002/ppp3.10182

Eidesen, P. B., Alsos, I. G., & Brochmann, C. (2015). Comparative analyses of plastid and AFLP data suggest different colonization history and asymmetric hybridization between Betula pubescens and B. nana. Molecular Ecology, 24, 3993–4009. https://doi.org/10.1111/mec.13289

Fowler, N. L., & Levin, D. A. (1984). Ecological constraints on the establishment of a novel polyploid in competition with its diploid progenitor. The American Naturalist, 124, 703–711. https://doi.org/10.1086/284307

Fujita, M. K., Leaché, A. D., Burbrink, F. T., McGuire, J. A., & Moritz, C. (2012). Coalescent-based species delimitation in an integrative taxonomy. Trends in ecology & evolution, 27, 480–488. https://doi.org/10.1016/j.tree.2012.04.012

Funk, W. C., Caminer, M., & Ron, S. R. (2012). High levels of cryptic species diversity uncovered in Amazonian frogs. Proceedings of the Royal Society B, 279, 1806–1814. https://doi.org/10.1098/rspb.2011.1653

Grewe, F., Huang, J. P., Leavitt, S. D., & Lumbsch, H. T. (2017). Reference-based RADseq resolves robust relationships among closely related species of lichen-forming fungi using metagenomic DNA. Scientific Reports, 7, 1–11. https://doi.org/10.1038/s41598-017-09906-7

Hall, T. A. 1999. BioEdit: a user-friendly biological sequence alignment editor and analysis program for Windows 95/98/NT. Nucleic Acids Symposium Series.

Hausdorf, B. (2011). Progress toward a general species concept. Evolution: International Journal of Organic Evolution, 65, 923–931. https://doi.org/10.1111/j.1558-5646.2011.01231.x

Hey, J. (2006). On the failure of modern species concepts. Trends in ecology & evolution, 21, 447–450. https://doi.org/10.1016/j.tree.2006.05.011

Hu, Y. N., Zhao, L., Buggs, R. J., Zhang, X. M., Li, J., & Wang, N. (2019). Population structure of Betula albosinensis and Betula platyphylla: evidence for hybridization and a cryptic lineage. Annals of Botany, 123, 1179–1189. https://doi.org/10.1093/aob/mcz024

Husband, B. C., & Sabara, H. A. (2004). Reproductive isolation between autotetraploids and their diploid progenitors in fireweed, Chamerion angustifolium (Onagraceae). New Phytologist, 161, 703–713. https://doi.org/10.1046/j.1469-8137.2004.00998.x

Jombart, T. (2008). adegenet: a R package for the multivariate analysis of genetic markers. Bioinformatics, 24, 1403–1405. https://doi.org/10.1093/bioinformatics/btn129

Levin, D. A. (1975). Minority cytotype exclusion in local plant populations. Taxon, 24, 35–43. https://doi.org/10.2307/1218997

Li, H. (2011). A statistical framework for SNP calling, mutation discovery, association mapping and population genetical parameter estimation from sequencing data. Bioinformatics, 27, 2987–2993. https://doi.org/10.1093/bioinformatics/btr509

Li, H., & Durbin, R. (2009). Fast and accurate short read alignment with Burrows–Wheeler transform. Bioinformatics, 25, 1754–1760. https://doi.org/10.1093/bioinformatics/btp324

Li, H., Handsaker, B., Wysoker, A., Fennell, T., Ruan, J., Homer, N., Marth, G., Abecasis, G., & Durbin, R. (2009). The sequence alignment/map format and SAMtools. Bioinformatics, 25, 2078–2079. https://doi.org/10.1093/bioinformatics/btp352

Li, P. C., & Skvortsov, A. K. (1999). Betulaceae. In: Wu ZY, Raven PH (eds) Flora of China, 4. Beijing: Science Press; St. Louis: Missouri Botanical Garden Press, 286–313.

Mayr, E. (1942). Systematics and the Origin of Species, From the Viewpoint of a Zoologist (No. 13). Cambridge, Massachusetts: Harvard University Press.

McKenna, A., Hanna, M., Banks, E., Sivachenko, A., Cibulskis, K., Kernytsky, A., et al. (2010). The Genome Analysis Toolkit: a MapReduce framework for analyzing next-generation DNA sequencing data. Genome Research, 20, 1297–1303. http://www.genome.org/cgi/doi/10.1101/gr.107524.110

Newton, L. G., Starrett, J., Hendrixson, B. E., Derkarabetian, S., & Bond, J. E. (2020). Integrative species delimitation reveals cryptic diversity in the southern Appalachian Antrodiaetus unicolor (Araneae: Antrodiaetidae) species complex. Molecular Ecology, 29, 2269–2287. https://doi.org/10.1111/mec.15483

Roccaforte, K., Russo, S. E., & Pilson, D. (2015). Hybridization and reproductive isolation between diploid Erythronium mesochoreum and its tetraploid congener E. albidum (Liliaceae). Evolution, 69, 1375–1389. https://doi.org/10.1111/evo.12666

Salojärvi, J., Smolander, O. P., Nieminen, K., Rajaraman, S., Safronov, O., Safdari, P., et al. (2017). Genome sequencing and population genomic analyses provide insights into the adaptive landscape of silver birch. Nature genetics, 49, 904–912. https://doi.org/10.1038/ng.3862

Skvortsov, A. K. (1997). Taxonomic notes on Betula. 1. section Betulaster. Harvard Papers in Botany, 65–70. https://www.jstor.org/stable/41761538

Smith, L. T., Magdalena, C., Przelomska, N. A., Pérez-Escobar, O. A., Melgar-Gómez, D. G., Beck, S., et al. (2022). Revised species delimitation in the giant water lily genus Victoria (Nymphaeaceae) confirms a new species and has implications for its conservation. Frontiers in Plant Science, 2241. https://doi.org/10.3389/fpls.2022.883151

Stamatakis, A. (2006). RAxML-VI-HPC: maximum likelihood-based phylogenetic analyses with thousands of taxa and mixed models. Bioinformatics, 22, 2688–2690. https://doi.org/10.1093/bioinformatics/btl446

Tarieiev, A., Olshanskyi, I., Gailing, O., & Krutovsky, K. V. (2019). Taxonomy of dark-and white-barked birches related to Betula pendula and B. pubescens (Betulaceae) in Ukraine based on both morphological traits and DNA markers. Botanical Journal of the Linnean Society, 191, 142–154. https://doi.org/10.1093/botlinnean/boz031

Thórsson, Æ. T., Pálsson, S., Lascoux, M., & Anamthawat-Jónsson, K. (2010). Introgression and phylogeography of Betula nana (diploid), B. pubescens (tetraploid) and their triploid hybrids in Iceland inferred from cpDNA haplotype variation. Journal of Biogeography, 37, 2098–2110. https://doi.org/10.1111/j.1365-2699.2010.02353.x

Tsuda, Y., Semerikov, V., Sebastiani, F., Vendramin, G. G., & Lascoux, M. (2017). Multispecies genetic structure and hybridization in the Betula genus across Eurasia. Molecular Ecology, 26, 589–605. https://doi.org/10.1111/mec.13885

Twyford, A. D., & Ennos, R. (2012). Next-generation hybridization and introgression. Heredity, 108, 179–189. https://doi.org/10.1038/hdy.2011.68

Wang, L. W., Ding, J. Y., Borrell, J. S., Cheek, M., McAllister, H. A., Wang, F. F., et al. (2022). Molecular and morphological analyses clarify species delimitation in section Costatae and reveal Betula buggsii sp. nov.(sect. Costatae, Betulaceae) in China. Annals of Botany, 129, 415–428. https://doi.org/10.1093/aob/mcac001

Wang, N., Borrell, J., & Buggs, R. (2014a). Is the Atkinson discriminant function a reliable method for distinguishing between Betula pendula and B. pubescens (Betulaceae)? New Journal of Botany, 4, 90–94. https://doi.org/10.1179/2042349714Y.0000000044

Wang, N., Borrell, J. S., Bodles, W. J., Kuttapitiya, A., Nichols, R. A., & Buggs, R. J. (2014b). Molecular footprints of the Holocene retreat of dwarf birch in Britain. Molecular Ecology, 23, 2771–2782. https://doi.org/10.1111/mec.12768

Wang, N., Kelly, L. J., McAllister, H. A., Zohren, J., & Buggs, R. J. (2021). Resolving phylogeny and polyploid parentage using genus-wide genome-wide sequence data from birch trees. Molecular Phylogenetics and Evolution, 160, 107126. https://doi.org/10.1016/j.ympev.2021.107126

Wang, N., McAllister, H. A., Bartlett, P. R., & Buggs, R. J. (2016). Molecular phylogeny and genome size evolution of the genus Betula (Betulaceae). Annals of Botany, 117, 1023–1035. https://doi.org/10.1093/aob/mcw048

Wang, N., Thomson, M., Bodles, W. J., Crawford, R. M., Hunt, H. V., Featherstone, A. W., Pellicer, J., & Buggs, R. J. (2013). Genome sequence of dwarf birch (Betula nana) and cross-species RAD markers. Molecular Ecology, 22, 3098–3111. https://doi.org/10.1111/mec.12131

White, T. J., Bruns, T., Lee, S., & Taylor, J. (1990). Amplification and direct sequencing of fungal ribosomal RNA genes for phylogenetics. PCR Protocols: A Guide to Methods and Applications, 18, 315–322. https://doi.org/10.1016/B978-0-12-372180-8.50042-1

Zeng, J., Li, J. H., & Chen, Z. D. (2008). A new species of Betula section Betulaster (Betulaceae) from China. Botanical Journal of the Linnean Society, 156, 523–528. https://doi.org/10.1111/j.1095-8339.2007.00764.x

Zeng, Z., Ren, B. Q., Zhu, J. Y., & Chen, Z. D. (2014). Betula hainanensis (Betulaster, Betulaceae), a new species from Hainan Island, China. Annales Botanici Fennici, 51, 399–402. http://dx.doi.org/10.5735/085.051.0606

Zhang, J. L., Zhang, C., Gao, L. M., Yang, J. B., & Li, H. T. (2007). Natural hybridization origin of Rhododendron agastum (Ericaceae) in Yunnan, China: inferred from morphological and molecular evidence. Journal of Plant Research, 120, 457–463. https://doi.org/10.1007/s10265-007-0076-1

Zohren, J., Wang, N., Kardailsky, I., Borrell, J. S., Joecker, A., Nichols, R. A., & Buggs, R. J. (2016). Unidirectional diploid-tetraploid introgression among British birch trees with shifting ranges shown by restriction site-associated markers. Molecular Ecology, 25, 2413–2426. https://doi.org/10.1111/mec.13644

