## Supplemental tables and figures for "A nearly 30-years living collection from the Royal Botanic Garden Edinburgh is a new species: a case study of *Betula mcallisterii* sp. nov. (sect. *Acuminatae*, Betulaceae) and its little hybridization with *Betula luminifera*"

**Supplementary data**

**Table S1** Detailed information on all samples used in the present study.

| Individuals | Species | Latitude  (°N) | Longitude  (°E) | Clean reads | Mapped reads | Mapped rate (%) | Variants |
| --- | --- | --- | --- | --- | --- | --- | --- |
| SZZ013 | *B. luminifera* | 25.79 | 105.15 | 15,408,170 | 14,251,142 | 92.49 | 1,335,748 |
| SZZ1 | *B. luminifera* | 25.79 | 105.15 | 19,326,092 | 17,331,575 | 89.68 | 1,573,336 |
| SZZ4 | *B. luminifera* | 25.79 | 105.15 | 14,925,864 | 12,969,418 | 86.89 | 1,481,376 |
| SZZ5 | *B. luminifera* | 25.79 | 105.15 | 16,725,832 | 14,405,742 | 86.13 | 1,456,353 |
| YLB001 | *B. luminifera* | 26.04 | 105.74 | 31,662,030 | 2,789,213 | 8.81 | 340,979 |
| XZ018 | *B. luminifera* | 26.28 | 107.43 | 21,255,620 | 9,323,543 | 43.86 | 1,262,284 |
| PD004 | *B. luminifera* | 27.23 | 105.86 | 18,219,898 | 6,122,000 | 33.60 | 1,074,434 |
| PD005 | *B. luminifera* | 27.23 | 105.86 | 15,754,046 | 13,448,460 | 85.37 | 1,340,913 |
| HJG019 | *B. luminifera* | 28.11 | 106.85 | 13,780,112 | 11,023,752 | 80.00 | 1,168,511 |
| TG007 | *B. luminifera* | 28.32 | 106.15 | 23,345,108 | 9,050,503 | 38.77 | 1,242,566 |
| YFZ003 | *B. luminifera* | 28.79 | 103.93 | 22,307,656 | 21,010,707 | 94.19 | 1,586,984 |
| YFZ011 | *B. luminifera* | 28.79 | 103.93 | 19,629,442 | 18,612,178 | 94.82 | 1,489,118 |
| YFZ017 | *B. luminifera* | 28.79 | 103.93 | 24,627,126 | 23,127,137 | 93.91 | 1,631,103 |
| YFZ023 | *B. luminifera* | 28.79 | 103.93 | 19,984,834 | 17,540,593 | 87.77 | 1,375,690 |
| YFZ025 | *B. luminifera* | 28.79 | 103.93 | 20,559,120 | 18,391,947 | 89.46 | 1,490,749 |
| WLP21-002 | *B. luminifera* | 31.35 | 109.90 | 16,577,240 | 13,139,252 | 79.26 | 1,251,325 |
| WLP21-007 | *B. luminifera* | 31.35 | 109.90 | 13,470,328 | 12,471,919 | 92.59 | 1,213,053 |
| WLP21-021 | *B. luminifera* | 31.35 | 109.90 | 22,883,706 | 15,261,146 | 66.69 | 1,300,982 |
| WLP21-023 | *B. luminifera* | 31.35 | 109.90 | 24,504,488 | 19,754,450 | 80.62 | 1,444,535 |
| WLP21-028 | *B. luminifera* | 31.35 | 109.90 | 19,522,140 | 14,535,074 | 74.45 | 1,312,265 |
| CKX18-028 | *B. luminifera* | 31.85 | 108.68 | 15,039,260 | 14,199,252 | 94.41 | 1,359,998 |
| CKX18-029 | *B. luminifera* | 31.85 | 108.68 | 18,695,080 | 16,346,646 | 87.44 | 1,668,411 |
| CKX18-031 | *B. luminifera* | 31.85 | 108.68 | 13,295,132 | 11,292,091 | 84.93 | 1,714,702 |
| CKX18-034 | *B. luminifera* | 31.85 | 108.68 | 14,437,658 | 13,441,155 | 93.10 | 1,382,358 |
| CKX18-035 | *B. luminifera* | 31.85 | 108.68 | 14,052,940 | 12,833,204 | 91.32 | 1,489,568 |
| CKX18-036 | *B. luminifera* | 31.85 | 108.68 | 17,231,644 | 15,771,033 | 91.52 | 1,401,013 |
| CKX18-040 | *B. luminifera* | 31.85 | 108.68 | 13,581,650 | 12,293,962 | 90.52 | 1,322,814 |
| DBS026 | *B. luminifera* | 31.86 | 109.13 | 16,758,432 | 15,290,270 | 91.24 | 1,211,286 |
| DBS036 | *B. luminifera* | 31.86 | 109.13 | 14,316,708 | 13,047,364 | 91.13 | 1,253,017 |
| EDL20-029 | *B. luminifera* | 33.38 | 106.56 | 14,737,340 | 13,188,110 | 89.49 | 1,070,443 |
| EDL20-033 | *B. luminifera* | 33.38 | 106.56 | 14,037,378 | 12,722,634 | 90.63 | 1,281,548 |
| EDL20-036 | *B. luminifera* | 33.38 | 106.56 | 16,739,198 | 15,565,052 | 92.99 | 1,356,499 |
| EDL20-039 | *B. luminifera* | 33.38 | 106.56 | 14,756,148 | 13,829,648 | 93.72 | 1,271,696 |
| EDL20-041 | *B. luminifera* | 33.38 | 106.56 | 19,233,980 | 12,910,878 | 67.13 | 1,217,200 |
| EDL20-045 | *B. luminifera* | 33.38 | 106.56 | 15,986,882 | 14,909,076 | 93.26 | 1,342,563 |
| EDL20-048 | *B. luminifera* | 33.38 | 106.56 | 14,081,002 | 12,597,227 | 89.46 | 1,264,841 |
| EDL20-051 | *B. luminifera* | 33.38 | 106.56 | 13,027,046 | 12,039,137 | 92.42 | 1,189,389 |
| EDL20-055 | *B. luminifera* | 33.38 | 106.56 | 13,108,438 | 12,202,004 | 93.09 | 1,243,450 |
| NSX20-001 | *B. luminifera* | 33.50 | 108.42 | 12,456,436 | 11,440,999 | 91.85 | 1,135,402 |
| NSX20-006 | *B. luminifera* | 33.50 | 108.42 | 13,833,224 | 12,792,080 | 92.47 | 1,193,730 |
| NSX20-009 | *B. luminifera* | 33.50 | 108.42 | 22,466,280 | 20,851,292 | 92.81 | 1,293,034 |
| NSX20-022 | *B. luminifera* | 33.50 | 108.42 | 17,597,920 | 16,388,975 | 93.13 | 1,391,904 |
| MXJ001 | *B. luminifera* | 33.51 | 106.65 | 16,521,680 | 15,459,277 | 93.57 | 1,361,714 |
| MXJ003 | *B. luminifera* | 33.51 | 106.65 | 18,007,306 | 16,852,362 | 93.59 | 1,226,451 |
| MXJ005 | *B. luminifera* | 33.51 | 106.65 | 20,351,920 | 15,079,954 | 74.10 | 1,161,717 |
| MXJ008 | *B. luminifera* | 33.51 | 106.65 | 19,891,046 | 18,513,609 | 93.08 | 1,297,920 |
| MXJ010 | *B. luminifera* | 33.51 | 106.65 | 16,295,246 | 14,959,510 | 91.80 | 1,154,496 |
| DBS023 | Unidentified | 31.86 | 109.13 | 18,470,430 | 17,308,390 | 93.71 | 838,796 |
| EDL20-003 | Unidentified | 33.38 | 106.56 | 16,566,072 | 15,489,626 | 93.50 | 824,157 |
| EDL20-006 | Unidentified | 33.38 | 106.56 | 16,633,850 | 15,786,411 | 94.91 | 989,800 |
| EDL20-009 | Unidentified | 33.38 | 106.56 | 18,657,068 | 17,702,009 | 94.88 | 907,014 |
| EDL20-014 | Unidentified | 33.38 | 106.56 | 17,040,046 | 15,192,931 | 89.16 | 1,283,571 |
| EDL20-017 | Unidentified | 33.38 | 106.56 | 14,098,230 | 13,308,248 | 94.40 | 979,525 |
| EDL20-022 | Unidentified | 33.38 | 106.56 | 15,843,810 | 14,731,193 | 92.98 | 1,248,972 |
| EDL20-026 | Unidentified | 33.38 | 106.56 |  |  |  |  |
| EDL20-032 | Unidentified | 33.38 | 106.56 | 15,435,784 | 13,926,691 | 90.22 | 755,230 |
| EDL20-035 | Unidentified | 33.38 | 106.56 | 13,612,162 | 12,820,783 | 94.19 | 940,785 |
| EDL20-037 | Unidentified | 33.38 | 106.56 | 15,210,572 | 14,136,421 | 92.94 | 903,784 |
| EDL20-046 | Unidentified | 33.38 | 106.56 | 14,843,744 | 14,043,880 | 94.61 | 853,885 |
| EDL20-052 | Unidentified | 33.38 | 106.56 | 13,566,732 | 12,789,635 | 94.27 | 905,426 |
| EDL20-054 | Unidentified | 33.38 | 106.56 | 12,956,516 | 12,258,724 | 94.61 | 916,454 |
| EDL20-056 | Unidentified | 33.38 | 106.56 | 15,331,162 | 14,510,529 | 94.65 | 932,930 |
| EDL21-004 | Unidentified | 33.38 | 106.56 |  |  |  |  |
| EDL21-012 | Unidentified | 33.38 | 106.56 | 26,951,064 | 13,781,877 | 51.14 | 859,687 |
| EDL21-017 | Unidentified | 33.38 | 106.56 |  |  |  |  |
| XYB20-001 | Unidentified | 33.47 | 108.50 |  |  |  |  |
| XYB20-004 | Unidentified | 33.47 | 108.50 |  |  |  |  |
| XYB21-001 | Unidentified | 33.47 | 108.50 | 29,327,080 | 27,650,130 | 94.28 | 1,317,184 |
| XYB21-002 | Unidentified | 33.47 | 108.50 | 31,562,674 | 29,246,050 | 92.66 | 1,280,401 |
| NSX19-007 | Unidentified | 33.50 | 108.42 |  |  |  |  |
| NSX19-009 | Unidentified | 33.50 | 108.42 |  |  |  |  |
| NSX19-012 | Unidentified | 33.50 | 108.42 | 17,780,376 | 16,667,704 | 93.74 | 3,667,744 |
| NSX19-022 | Unidentified | 33.50 | 108.42 | 13,145,524 | 12,472,356 | 94.88 | 859,973 |
| NSX20-015 | Unidentified | 33.50 | 108.42 | 12,276,032 | 11,494,207 | 93.63 | 755,755 |
| NSX20-017 | Unidentified | 33.50 | 108.42 | 13,731,578 | 12,997,801 | 94.66 | 853,345 |
| NSX20-020 | Unidentified | 33.50 | 108.42 | 14,636,686 | 13,815,501 | 94.39 | 793,072 |
| NSX20-027 | Unidentified | 33.50 | 108.42 | 19,510,274 | 18,285,370 | 93.72 | 868,746 |
| NSX20-030 | Unidentified | 33.50 | 108.42 | 20,032,244 | 18,846,778 | 94.08 | 942,224 |
| NSX20-034 | Unidentified | 33.50 | 108.42 | 15,742,126 | 14,918,637 | 94.77 | 800,567 |
| NSX20-037 | Unidentified | 33.50 | 108.42 | 17,196,280 | 16,296,191 | 94.77 | 858,882 |
| NSX20-040 | Unidentified | 33.50 | 108.42 | 16,878,752 | 15,926,965 | 94.36 | 874,095 |
| NSX20-043 | Unidentified | 33.50 | 108.42 | 18,332,120 | 17,434,357 | 95.10 | 860,224 |
| NSX20-044 | Unidentified | 33.50 | 108.42 | 17,207,740 | 16,314,835 | 94.81 | 883,980 |
| NSX20-047 | Unidentified | 33.50 | 108.42 | 19,605,296 | 14,004,724 | 71.43 | 942,516 |
| NSX20-051 | Unidentified | 33.50 | 108.42 | 23,931,550 | 22,182,907 | 92.69 | 937,807 |
| NSX20-053 | Unidentified | 33.50 | 108.42 | 12,895,952 | 12,174,301 | 94.40 | 982,578 |
| NSX20-060 | Unidentified | 33.50 | 108.42 | 12,920,410 | 12,215,698 | 94.55 | 812,155 |
| NSX20-072 | Unidentified | 33.50 | 108.42 | 16,922,580 | 15,784,852 | 93.28 | 941,071 |
| NSX20-073 | Unidentified | 33.50 | 108.42 | 17,435,630 | 16,394,245 | 94.03 | 1,070,949 |
| NSX21-001 | Unidentified | 33.50 | 108.42 | 19,480,110 | 18,288,096 | 93.88 | 995,297 |
| NSX21-006 | Unidentified | 33.50 | 108.42 |  |  |  |  |
| NSX21-007 | Unidentified | 33.50 | 108.42 |  |  |  |  |
| NSX21-010 | Unidentified | 33.50 | 108.42 | 13,393,748 | 12,598,298 | 94.06 | 849,626 |
| NSX21-011 | Unidentified | 33.50 | 108.42 | 12,376,610 | 10,491,929 | 84.77 | 858,424 |
| NSX21-013 | Unidentified | 33.50 | 108.42 |  |  |  |  |
| NSX21-015 | Unidentified | 33.50 | 108.42 | 19,771,890 | 18,470,913 | 93.42 | 1,011,788 |
| NSX21-018 | Unidentified | 33.50 | 108.42 |  |  |  |  |
| NSX21-022 | Unidentified | 33.50 | 108.42 |  |  |  |  |
| NSX21-030 | Unidentified | 33.50 | 108.42 |  |  |  |  |

**Figure S1** Admixture results at K values from 2 to 6 based on 40,209 SNPs.

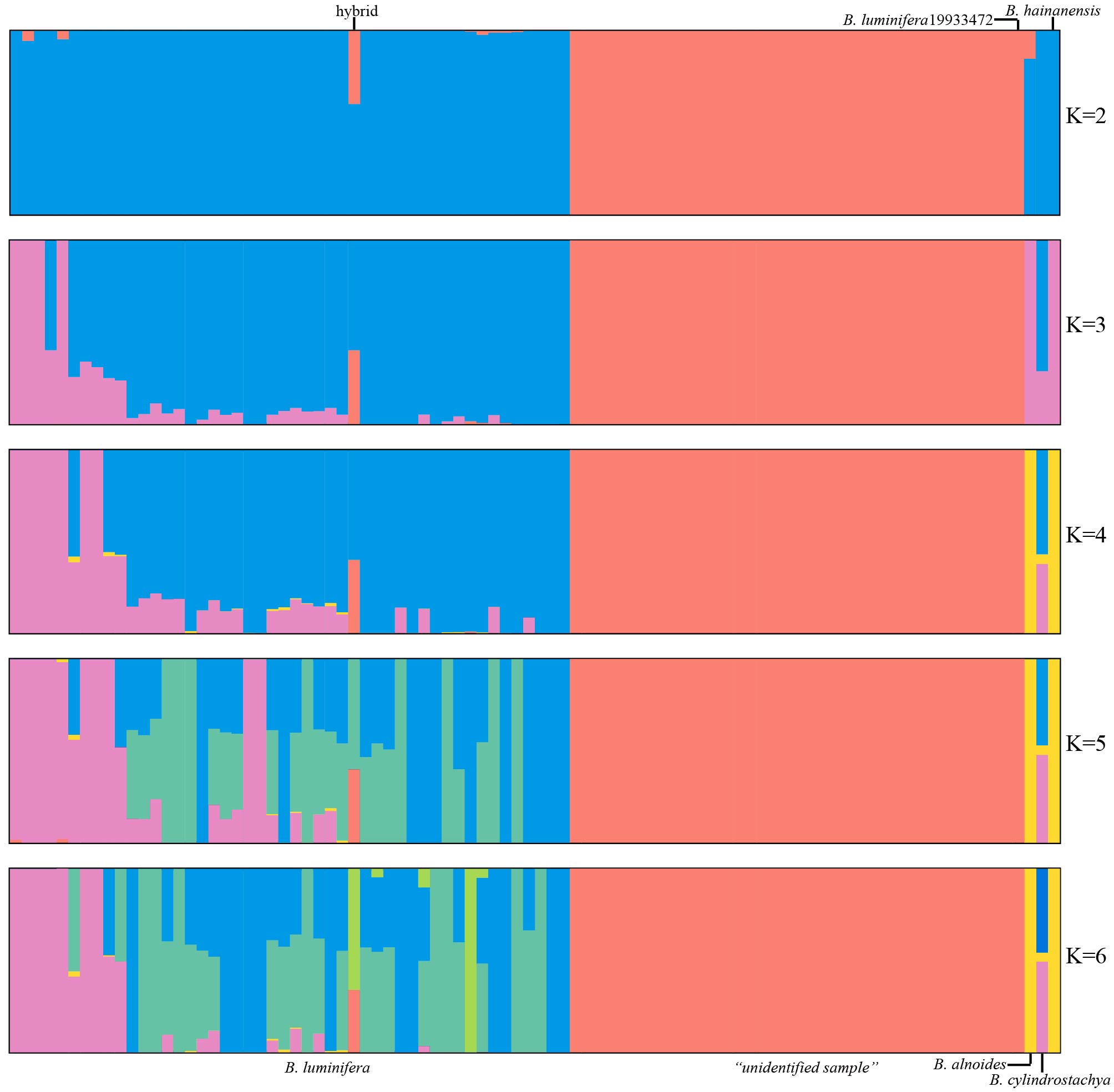

**Figure S2** Genome size estimation of the three *B. mcallisterii* samples. Red and blue peaks represent *B. mcallisterii* and the internal standard (tomato), respectively.

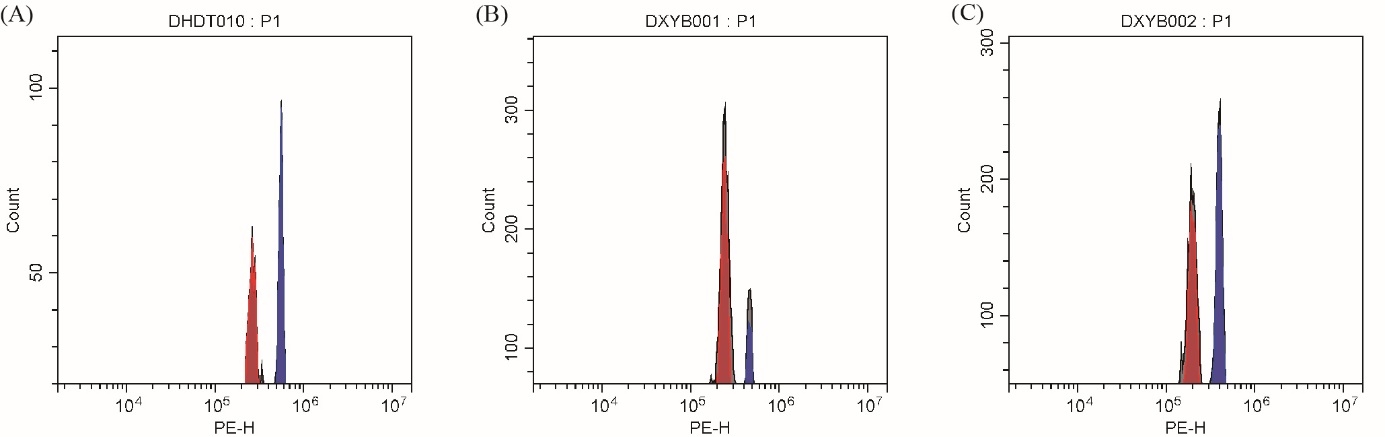

**Figure S3** Plots of read count ratios for heterozygous sites covered by at least 30 reads for *B. luminifera*.

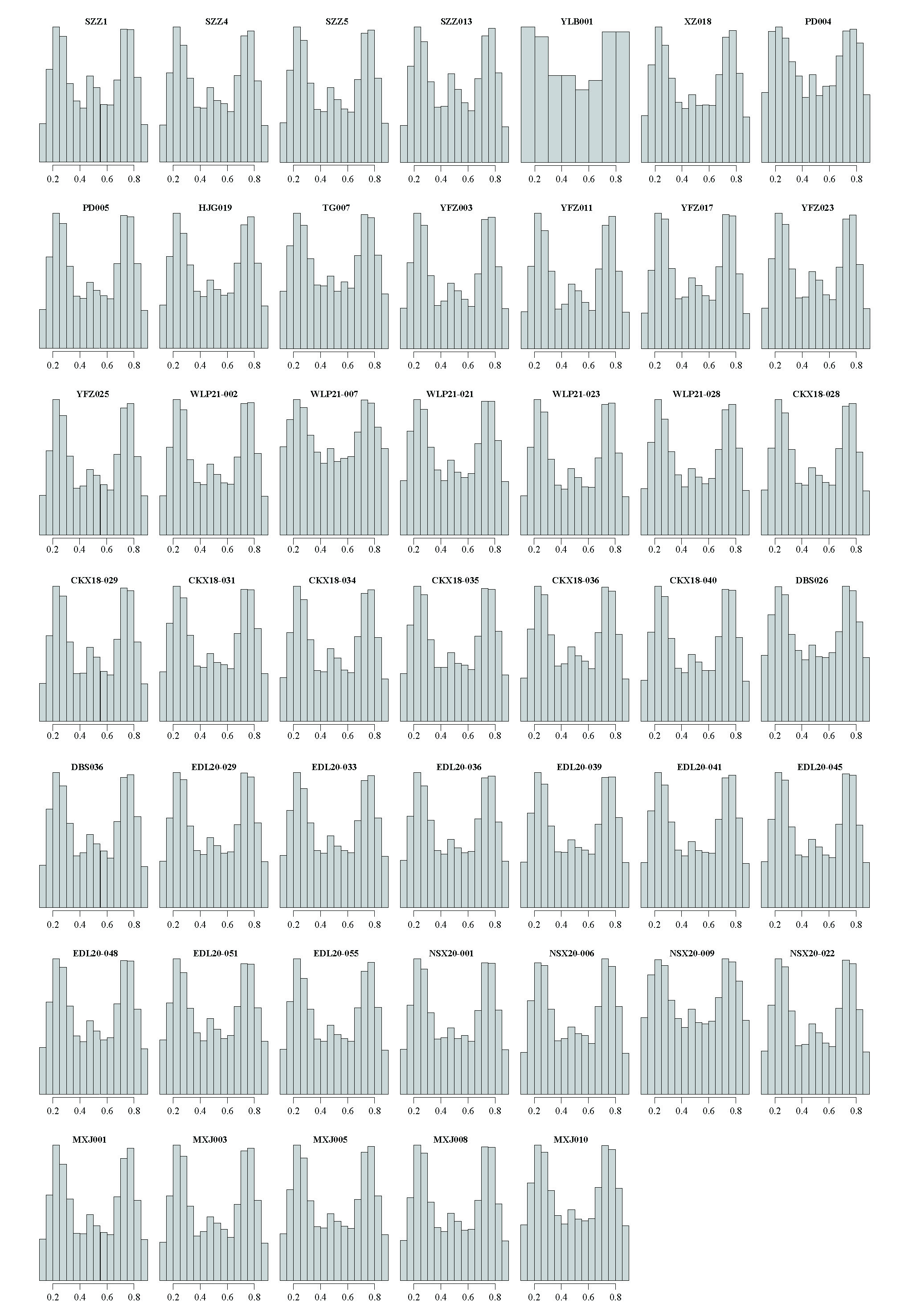

**Figure S4** Plots of read count ratios for heterozygous sites covered by at least 30 reads for the “unidentified sample”.

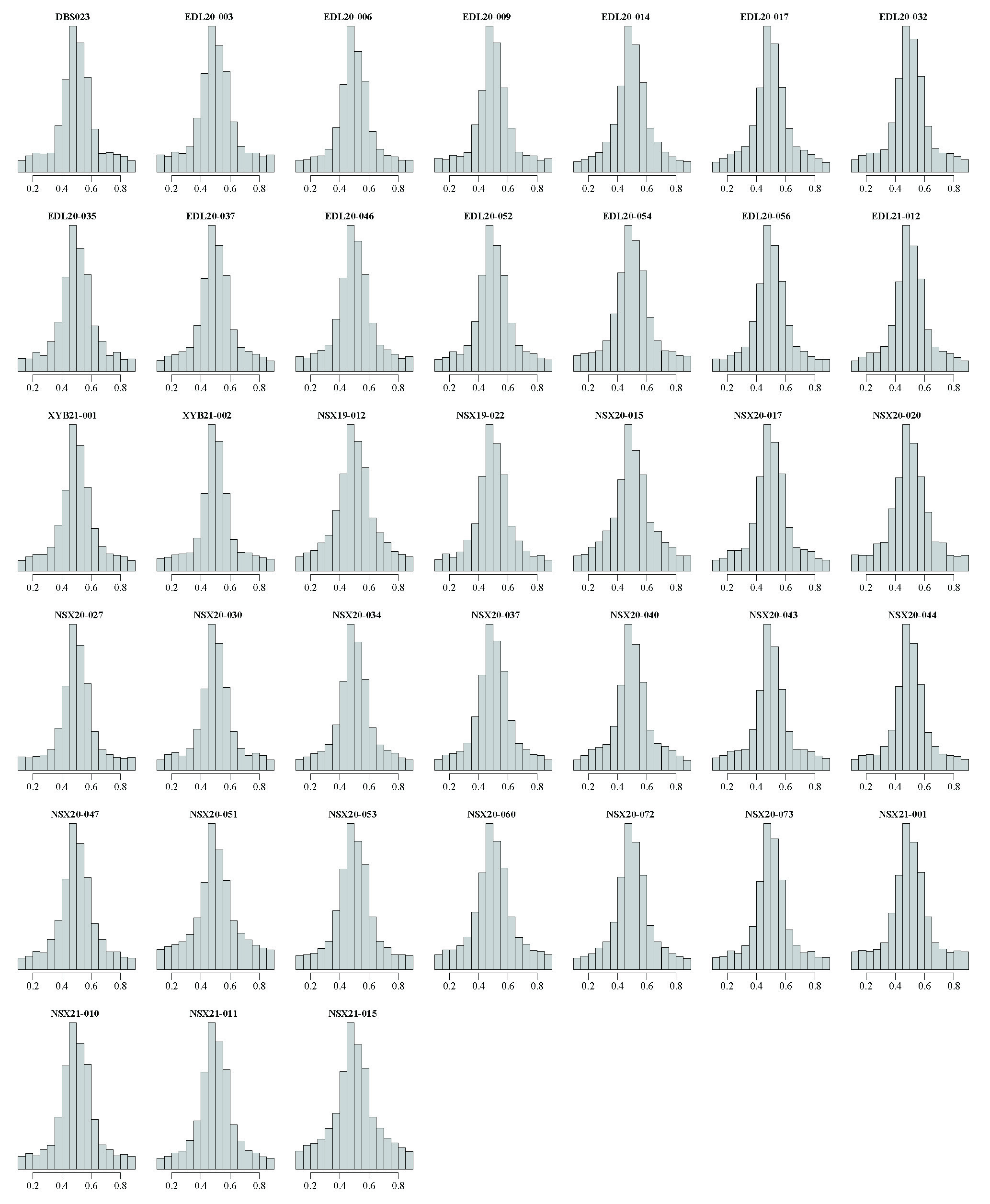

**Figure S5** Phylogenetic tree from the maximum-likelihood analysis of *B. mcallisterii* using ITS sequences.

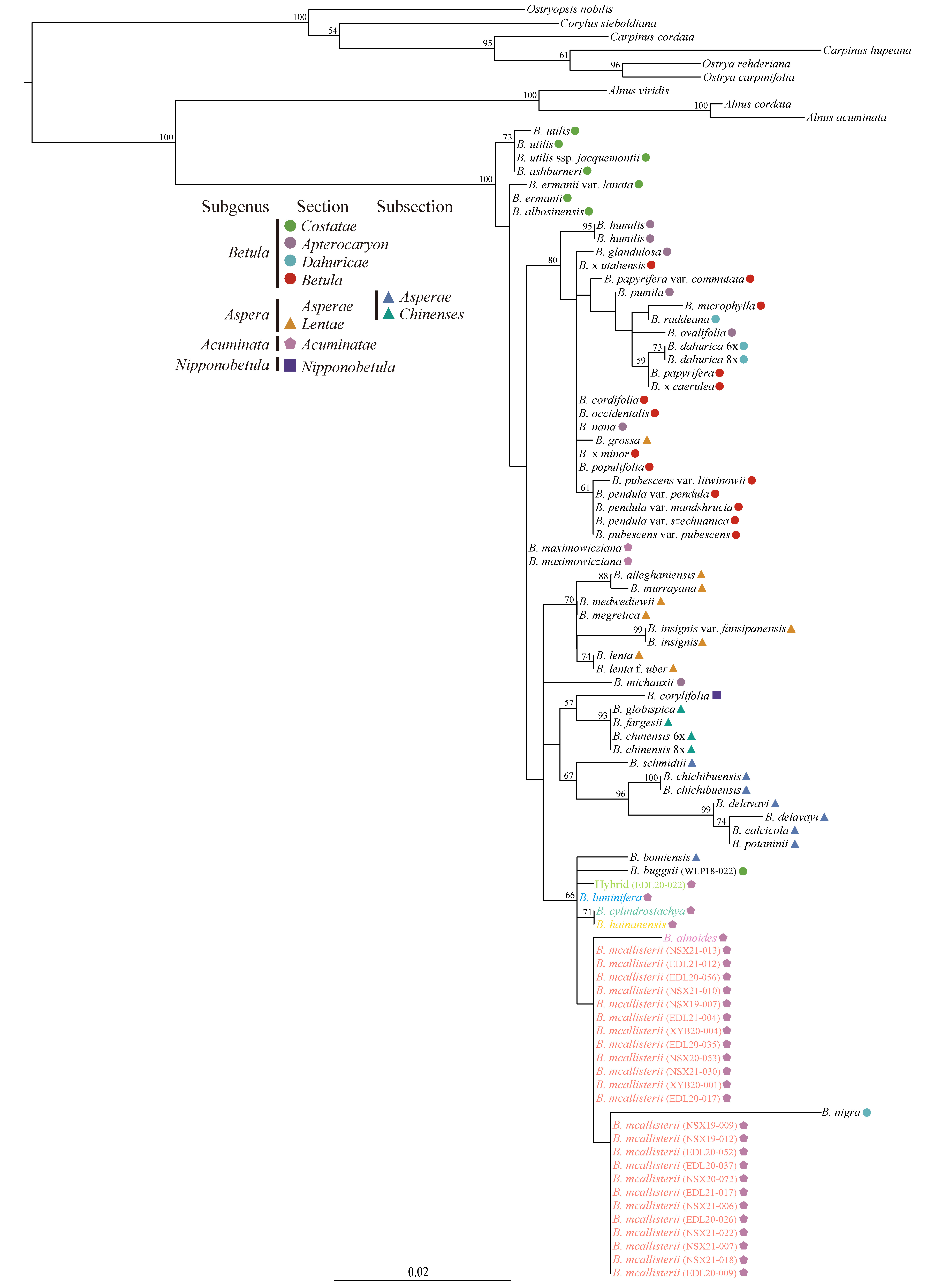

**Figure S6** Pictures of *Betula luminifera* 19933472 from the Royal Botanic Garden Edinburgh. (A) fruit; (B) male catkins; (C) and (D) bark.

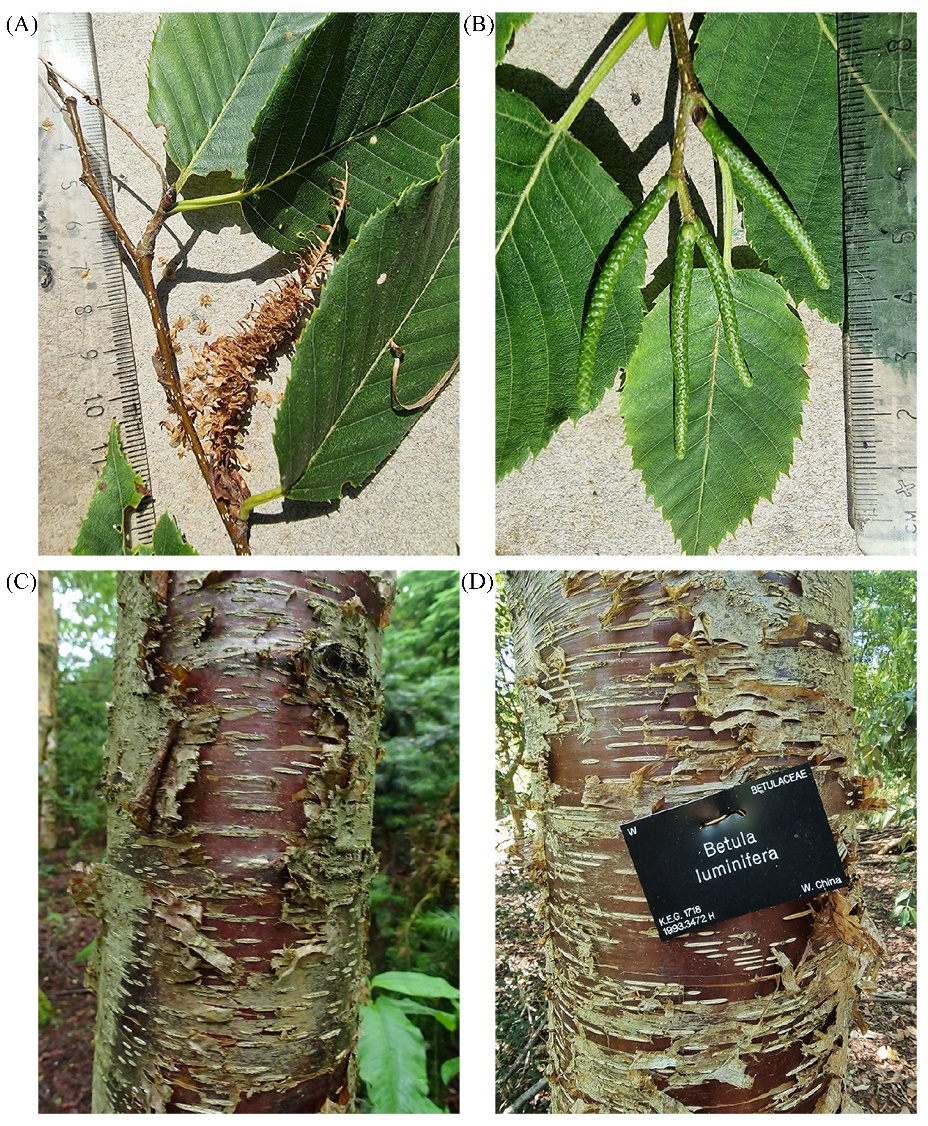
